# Revisiting the evolutionary analysis of mammalian CRISPs reveals positive selection

**DOI:** 10.1101/266940

**Authors:** Alberto Vicens, Claudia Treviño

## Abstract

Cysteine-rich secretory proteins (CRISPs) constitute a versatile family, with functions that include being components of reptilian venom and participation in mammalian reproduction. While non-mammalian vertebrates express a single CRISP gene, mammals generally express three CRISP paralogs. A previous study assessing the molecular evolution of vertebrate CRISPs revealed strong positive selection in reptilian CRISP and negative selection in mammalian CRISPs. In this study, we re-assessed molecular adaptation of mammalian CRISPs through an analysis of larger sequence datasets that represent mammalian diversity. Our analyses show evidence of recent episodes of positive selection for all mammalian CRISPs. Intensity of positive selection was heterogeneous both among CRISP paralogs (being stronger in CRISP3 than in CRISP1 and CRISP2) and across functional domains (having more impact on CRD or PR-1 domain). Analysis of episodic selection did not yield strong signatures of adaptive evolution in any particular mammalian group, suggesting that positive selection was more pervasive on mammalian CRISPs. Our findings provide evidence of adaptive evolution in a family of reproduction-related proteins, and offer interesting insights regarding the role of mammalian CRISPs in fertility and speciation.

## Introduction

CRISPs constitute a family of cysteine-rich secretory proteins with molecular weights of 20-30 kDa, characterized by the conservation of 16 cysteines that form disulfide bonds. CRISPs are considered dual function proteins, as they are composed by two different functional domains: an N-terminal CAP domain, known as plant pathogenesis-related 1 (PR-1), and a C-terminal region that comprises a hinge region and the characteristic cysteine-rich domain (CRD), which contains 10 of the 16 conserved cysteine residues (Gibbs et al. 2008).

CRISPs are exclusively found in vertebrates (Gibbs et al. 2008) and participate in different biological processes, depending on the context of expression and the specific taxonomic group. Non-mammalian vertebrates contain a single CRISP gene, and its biological function has been widely studied in reptiles, given that reptilian CRISPs are venom components (Yamazaki and Morita 2004; Gibbs and O’Bryan 2007). Most mammals have three CRISP paralogues (namely CRISP1, CRISP2 and CRISP3), although in mouse a fourth CRISP has been identified (Jalkanen et al. 2005; Gibbs et al. 2008). All mammalian CRISPs show enriched expression in the male reproductive tract, and they participate in different events of reproduction (Gibbs et al. 2008; Da Ros et al. 2015). CRISP1 is prominently expressed in the epididymis and binds to sperm during maturation, where it functions as sperm decapacitation factor and participates in both sperm-zona pellucida binding and gamete fusion (Cohen et al. 2001; Roberts et al. 2003; Cohen et al. 2011; Maldera et al. 2014). CRISP2 is a testicular protein that participates altogether with CRISP1 in sperm-egg fusion through binding to the same complementary site in the oocyte surface (Busso et al. 2007). CRISP3 shows a more widespread expression, but in humans it is also expressed in the epididymis, and it is found in ejaculated spermatozoa (Udby et al. 2005; Da Ros et al. 2015). Nonetheless, the role of CRISP3 in fertilization remains unclear.

Given that CRISP is a recently emerged protein family that has evolved in mammals to modulate multiple aspects of male reproduction, its members represent potential candidates as proteins that have diversified to enhance the fertilization success, particularly in competitive scenarios of post-copulatory sexual selection (Birkhead et al. 2008). In addition, the finding that mammalian CRISPs display different functions among closely related species (Da Ros et al. 2015) suggests that CRISPs could have undergone recent episodes of positive selection driving functional shifts. Therefore, an evolutionary study analyzing the molecular mechanisms along with the functional divergence of mammalian CRISPs, may give valuable insights into the biological activities of these reproduction-related proteins. In a previous study, Sunagar et al. (2012) evaluated the molecular evolution of reptilian and mammalian CRISPs. This study revealed that positive selection is strong in the evolution of reptilian CRISPs (particularly in snakes), whereas mammalian CRISPs are constrained by strong negative selection (Sunagar et al. 2012). However, this evolutionary analysis only compared a few mammalian lineages, and many of the sequences used in it were multiple transcript isoforms from the same species. These two factors can compromise the power of methods used to detect positive selection (Anisimova et al. 2001).

In the present study, we re-visited the selective pressures driving the evolution of the CRISP family using an interspecies approach in mammals. We first evaluated whether mammalian CRISPs underwent accelerated evolution following gene duplications. We then assessed the selective pressures driving the late evolution of mammalian CRISPs, and examined the targets of positive selection using structural and functional information. Finally, we explored whether episodes of adaptive evolution of CRISPs were associated with any particular mammalian groups.

## Results and Discussion

### Lack of positive selection in ancestral CRISPs

Mammalian CRISPs have different functions than their reptilian orthologs. Whereas reptilian CRISPs are venom compounds produced in salivary glands, mammalian CRISPs are non-toxicoferan proteins with prominent expression in male reproductive tissues and important roles in fertility. To identify potential adaptive changes that could lead to functional divergence between mammalian and reptilian CRISPs, we tested for positive selection in the ancestral branch of mammalian CRISP family. The selective constraints of protein-coding genes are evaluated from the estimation of the ratio of nonsynonymous substitutions (dN) to synonymous substitutions (dS). The dN/dS ratio is known as omega (ω), and the value of ω is indicative of the selective profile of a coding-gene. If ω < 1 the sequence is conserved by purifying selection, if ω = 1 the gene is neutrally evolving, and ω > 1 is indicative of positive selection. Applying likelihood branch-site models, the selection model detected 21.6 % of sites with ω > 1 (ω_f_ = 2.48) in the ancestral branch, but the Likelihood Ratio Test (LRT) was not significant as this model was compared to the null model constraining ω_f_ to 1 (LRT = 0, p = 1) (Fig. 1). This result suggests that there were not positive selection-driven changes in the divergence between reptilian and mammalian CRISP.

**Figure 1.**
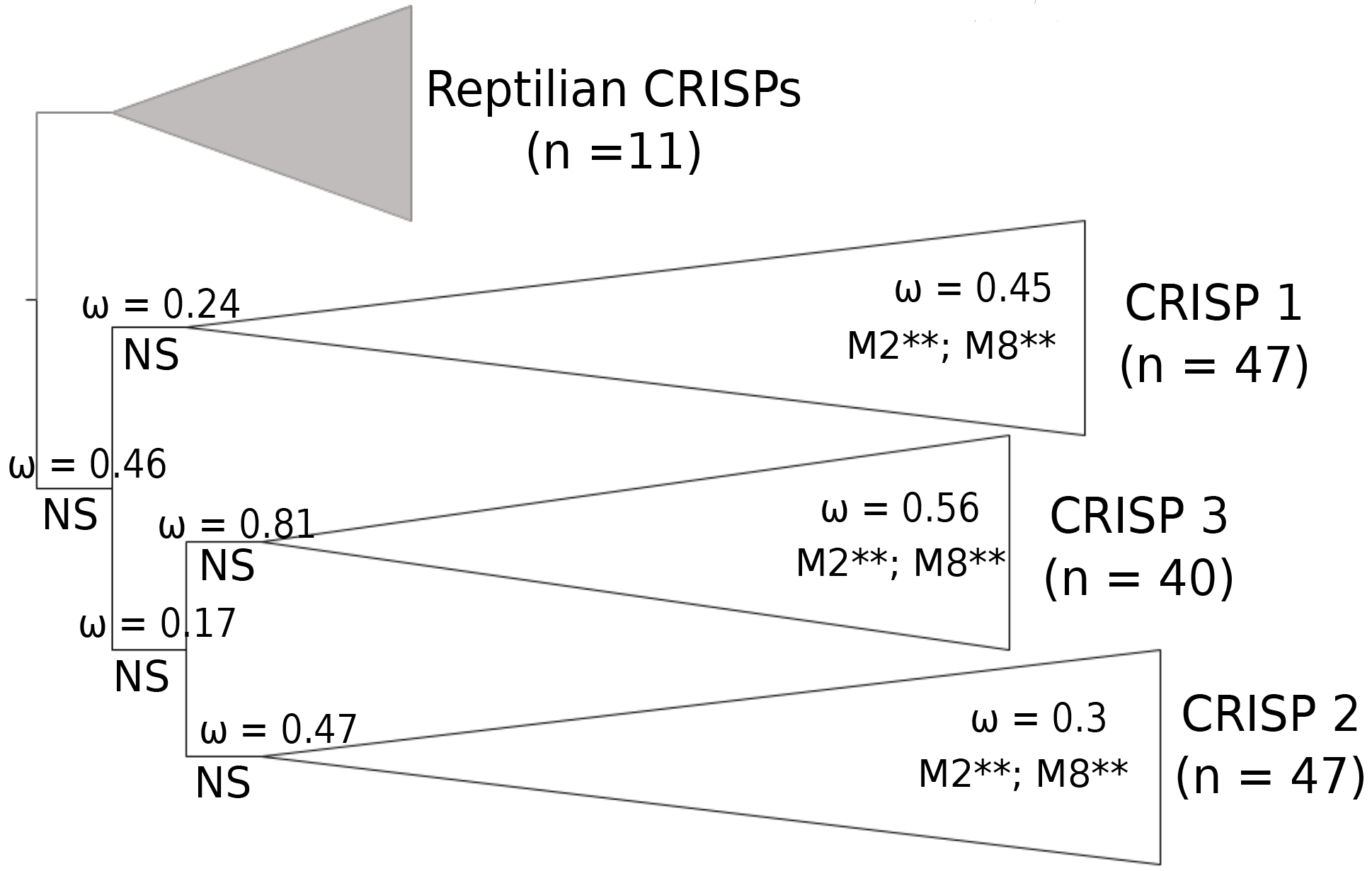
Selection analysis of CRISP family. The phylogenetic tree of CRISP family is showed with branches of each subfamily collapsed. Reptilian CRISPs were used as outgroup branches to analyze the ancestral state of mammalian CRISP. Branch-specific ω ratios estimated from two-ratio models are represented above internal branche. Results of likelihood ratio tests comparing selection and null branch-site models are indicated below branches (NS: no significant). In the terminal clades, overall ω estimates (M0) and significance of the best-fit site model is represented for each mammalian CRISP. The number of sequences retrieved of each CRISP is showed below protein names.

It has been well documented that episodes of positive selection often occur immediately following gene duplication, leading to functional divergence between daughter genes (Bielawski and Yang 2003; Zhang 2003). Therefore, we explored whether adaptive evolution occurred after duplication events that gave rise to the mammalian CRISP family. We first applied nested branch models to detect acceleration of evolutionary rate (i.e. increasing dN and ω) in the branches immediately following gene duplication. For the duplication that gave rise to the CRISP1 gene and the CRISP2/CRISP3 subfamily, a two-ratio model assuming one ω ratio for the branches separating the paralogs (ω_1_), and a background ω ratio for all other branches (ω_0_), was the best-fit model (LRT = 9.872; df = 1; p = 0.0016) (Table 1). Under this model, the evolutionary rate estimated for post-duplication branches (ω_1_ = 0.206) was lower than the background rate (ω_0_ = 0.464), suggesting strong purifying selection following the first duplication of the mammalian CRISP. A three-ratio model estimating different ω for the ancestral branches of each subfamily was not significant when compared to the two-ratio model (LRT = 1.676; df = 1; p = 0.195) (Table 1). For the duplication giving rise to CRISP2 and CRISP3, the LRT comparing the one- vs. the two-ratio model was not significant (LRT = 1.284; df = 1; p = 0.257), although parameter estimates suggested an acceleration of evolutionary rate following gene duplication (ω_1_ = 0.809), relative to the background rate (ω_0_ = 0.44) (Table 1). The three-ratio model showed identical likelihood to the two-ratio model, but the ancestral CRISP3 showed a higher evolutionary rate (ω_1_ = 0.81) (Table 1, Fig. 2). To further investigate whether episodes of positive selection occurred following gene duplication, we applied branch-site models. In this analysis, selection models were not significant respect to null models for any ancestral branch of the CRISP family (Table 2, Fig. 2), suggesting mammalian CRISPs did not undergo adaptive changes following gene duplication. Nonetheless, we must be cautious while interpreting these results, as branch-site tests can be conservative in detecting evidence of positive selection (Zhang et al. 2005).

**Table 1.** Parameter estimates and likelihood scores of nested branch-models in gene duplications of mammalian CRISP family.

| **Foreground branches** | **Model** | **np** | **Parameters** | **Likelihood** |
| --- | --- | --- | --- | --- |
| Duplication<br>CRISP1-CRISP2/CRISP3 | One-ratio (M0) | 306 | $\omega_0 = 0.457$ | -31977.845 |
|  | **Two-ratio** | **307** | **$\omega_0 = 0.464$**<br>**$\omega_1 = 0.206$** | **-31972.909** |
| | Three-ratio | 308 | $\omega_0 = 0.464$<br>$\omega_1 = 0.243$<br>$\omega_2 = 0.169$ | -31972.071 |
| Duplication<br>CRISP2-CRISP3 | One-ratio (M0) | 284 | $\omega_0 = 0.441$ | -28010.177 |
|  | **Two-ratio** | **285** | **$\omega_0 = 0.441$**<br>**$\omega_1 = 0.809$** | **-28009.535** |
| | Three-ratio | 286 | $\omega_0 = 0.441$<br>$\omega_1 = 0.809$<br>$\omega_2 = 0.473$ | -28009.535 |

**Figure 2.**
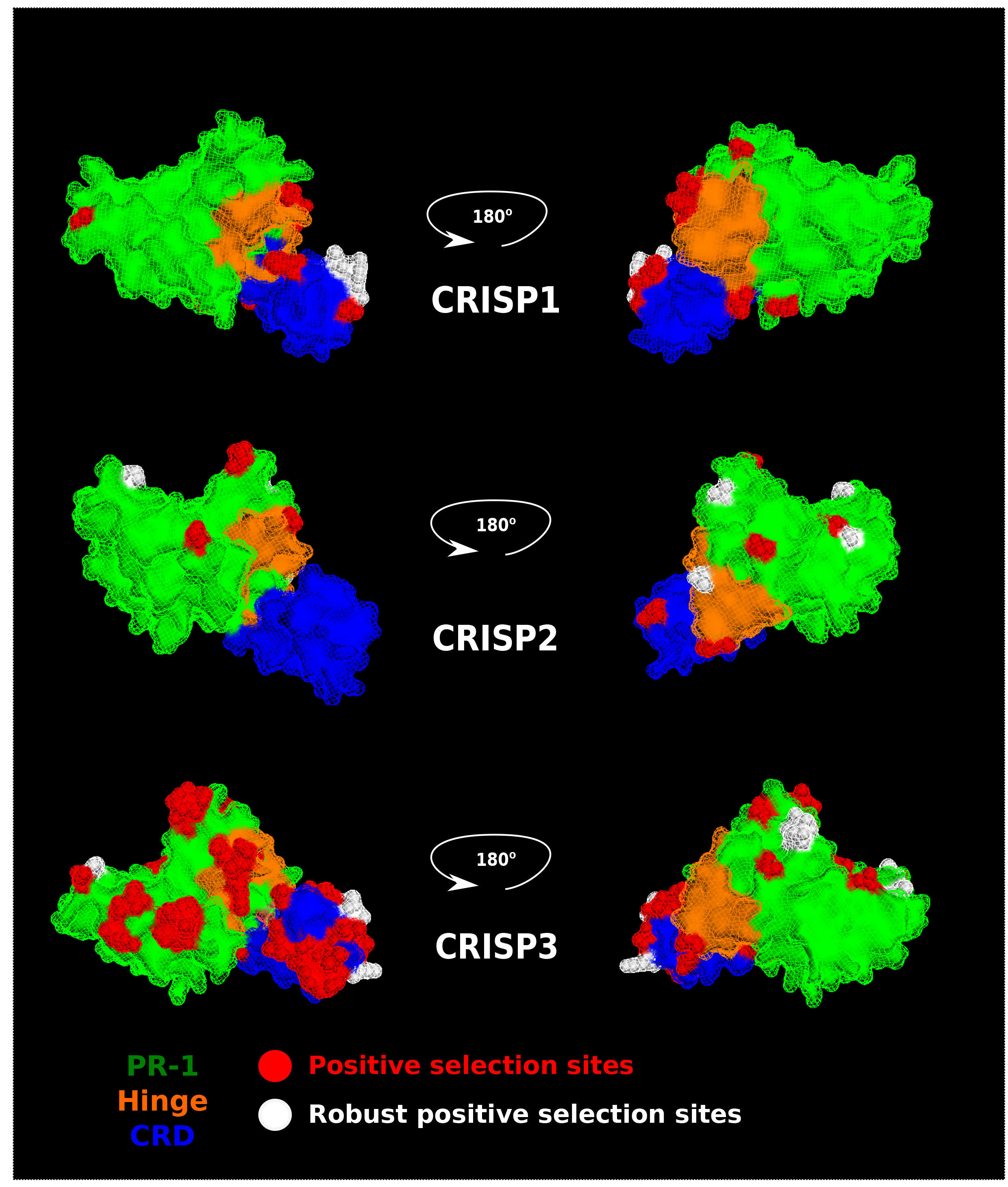
Three-dimensional models of hCRISP1, hCRISP2 and hCRISP3 are represented. Positively selected sites (PP > 0.95) detected by at least one selection model are indicated in red, and those detected by at least 3 selection models (robust) are indicated in white. Different domains of CRISPs are represented with different colors. Residue identified under of positive selection are indicated in Figures S1-S3.

**Table 2.** Parameter estimates and likelihood scores of site-specific models in mammalian CRISP

| Gene | Model | lnL | np | Parameters | PSS |
| --- | --- | --- | --- | --- | --- |
| CRISP1 | M0 | -9996.25 | 92 | $\omega_0 = 0.448$ | |
| | M1 | -9618.83 | 93 | $p_0 = 0.5, \omega_0 = 0.103$<br>$p_1 = 0.5, \omega_1 = 1$ | Not allowed |
| | M2 | -9608.24 | 95 | $p_0 = 0.5, \omega_0 = 0.108$<br>$p_1 = 0.42, \omega_1 = 1$<br>$p_2 = 0.5, \omega_2 = 1.87$ | 2 sites |
| | M7 | -9586.68 | 93 | $p = 0.35, q = 0.425$ | Not allowed |
| | M8a | -9582.83 | 94 | $p_0 = 0.702$<br>$p_1 = 0.297, \omega_1 = 1$<br>$p = 0.52, q = 1.737$ | Not allowed |
| | M8 | -9575.99 | 95 | $p_0 = 0.832$<br>$p_1 = 0.168, \omega_1 = 1.386$<br>$p = 0.43, q = 0.8$ | 11 sites |
| CRISP2 | M0 | -8030.23 | 98 | $\omega_0 = 0.303$ | |
| | M1 | -7692.13 | 101 | $p_0 = 0.64, \omega_0 = 0.069$<br>$p_1 = 0.356, \omega_1 = 1$ | Not allowed |
| | M2 | -7678.1359 | 103 | $p_0 = 0.62, \omega_0 = 0.067$<br>$p_1 = 0.327, \omega_1 = 1$<br>$p_2 = 0.05, \omega_2 = 2.26$ | 4 sites |
| | M7 | -7640.92 | 101 | $p = 0.35, q = 0.425$ | Not allowed |
| | M8a | -7639.34 | 102 | $p_0 = 0.853$<br>$p_1 = 0.147, \omega_1 = 1$<br>$p = 0.296, q = 1.419$ | Not allowed |
| | M8 | -7629.48 | 103 | $p_0 = 0.943$<br>$p_1 = 0.057, \omega_1 = 1.748$<br>$p = 0.257, q = 0.755$ | 7 sites |
| CRISP3<sup>a</sup> | M0 | -8580.89 | 76 | $\omega_0 = 0.513$ | |
| | M1 | -8137.5 | 77 | $p_0 = 0.541, \omega_0 = 0.091$<br>$p_1 = 0.46, \omega_1 = 1$ | Not allowed |
| | M2 | -8110.46 | 78 | $p_0 = 0.503, \omega_0 = 0.093$<br>$p_1 = 0.303, \omega_1 = 1$<br>$p_2 = 0.193, \omega_2 = 2.05$ | 10 sites |
| | M7 | -8103.50 | 77 | $p = 0.213, q = 0.25$ | Not allowed |
| | M8a | -8097.431 | 78 | $p_0 = 0.635$<br>$p_1 = 0.364, \omega_1 = 1$<br>$p = 0.39, q = 2.27$ | Not allowed |
| | M8 | -8075.39 | 79 | $p_0 = 0.747$<br>$p_1 = 0.253, \omega_1 = 1.669$<br>$p = 0.284, q = 0.698$ | 29 sites |

lnL: log likelihood estimated
np: number of parameters
<sup>a</sup>CRISP3-like copies of rodents with four CRISPs were discarded for CRISP3 analysis.

### Pervasive positive selection of mammalian CRISPs during late evolution

We first estimated an average evolutionary rate for each CRISP alignment. The overall estimates were ω = 0.448 for CRISP1, ω = 0.303 for CRISP2 and ω = 0.513 for CRISP3 (Table 2, Fig. 1). These global ω values were very close to those estimated in the study of Sunagar et al. (2012). While it is true that ω < 1 is indicative of purifying selection, global ω is a very conservative approach to infer selective pressures, as it averages over all sites of a sequence alignment and lineages of a phylogenetic tree. In any case, the observation that CRISP3 yielded higher global ω estimates is surprising, since this gene is the most widely expressed of the CRISP family (Gibbs et al. 2008).

To identify signatures of positive selection driving the divergence of mammalian CRISPs, we applied likelihood site-specific models. LRTs comparing selection (M2a, M8) with null (M1a, M7, M8a) models were significant for the three mammalian CRISPs (Table 2, Fig. 1, Table S2). To corroborate the presence of positive selection in mammalian CRISPs, we performed additional site selection methods that account for variation in dS rate among sites: fixed effects likelihood (FEL), random effects likelihood (REL) and likelihood-based counting (SLAC) methods. These approaches detected sites under positive selection with a PP > 0.95 in the three mammalian CRISPs (Table S3). The number of positively selected sites (PSS) was higher in CRISP3 under all site approaches except SLAC, supporting a stronger positive selection on this more widely expressed CRISP member. Our evolutionary models yielded a stronger signal of positive selection than the analysis previously applied by Sunagar et al. (2012) on mammalian CRISPs. The main difference between both studies is the number and the divergence of sequences employed. Sunagar et al. used datasets of between 10-22 sequences, many of them being identical isoforms from the same species originated by alternative splicing (Sunagar et al. 2012). Since the power of LRTs to detect positive selection decreases with few and more similar sequences (Anisimova et al. 2001), it is possible that the LRTs performed by Sunagar et al. (2012) did not have enough power to identify positive selection on mammalian CRISPs. In our analysis, we included more divergent sequences, increasing sharply the power of LRTs (Anisimova et al. 2001) Therefore, our analyses were more powerful in identifying positive selection, highlighting the importance of using large numbers and divergent sequences in studies of molecular adaptation.

To explore the functional implications of adaptive changes in mammalian CRISPs, we mapped PSS on three-dimensional structural models of human CRISPs (Fig. 2). In CRISP1, 6 of 13 residues under positive selection were located on CRD, covering 18.1% (6 of 37 amino acids) of this domain, associated with ion channel regulatory activity. Three and four sites under selection were detected in the PR-1 domain and in the hinge region, respectively. Meanwhile, positive selection was mainly focused on the PR-1 domain in CRISP2, with 8 of 13 PSS located in this functional region (Fig. 2). Three PSS mapped to the hinge and a single residue to the CRD. In spite of the strong signal of positive selection detected in the N-terminal region of CRISP2, no PSS mapped in the fusigenic signatures involved in gamete fusion (Ellerman et al. 2006), supporting the evolutionary conservation of these peptides. In CRISP3, a high number of PSS (25 of 46; 54.3 %) mapped to the PR-1 domain, and positive selection was particularly intense on the CRD domain, where 15 of the 38 residues defining such domain (39.4 %) were identified as PSS (Fig. 2). The location of PSS -in the primary structure of CRISPs- is shown in the Figures S1-S3. Since mammalian CRISPs have been described as modulators of sperm ion channels (Gibbs et al. 2006; Ernesto et al. 2015), it is possible that adaptive mutations, particularly in the CRD domain, could enhance or generate new interactions of CRISPs with ion transporters. On the other hand, mammalian CRISPs also participate in gamete interaction through binding to egg complementary sites (Cohen et al. 2011; Maldera et al. 2014), and thus adaptive mutations in CRISPs could facilitate sperm-egg interactions, thus contributing to reproductive success. Although it is beyond the scope of our study, future biochemical studies will be necessary to evaluate whether mutations in mammalian CRISPs cause adaptive phenotypic effects, and whether such changes correlate with reproductive fitness.

### Weak signal of episodic positive selection on mammalian lineages

Next, we assessed the selective pressures of CRISPs on the three major groups of mammals: Glires, Primates and Laurasiatheria. For this purpose, we applied two-ratio and branch-site models of selection (Yang and Nielsen 2002; Zhang et al. 2005). In CRISP1, two-ratio models estimated a lower foreground ω for Glires, suggesting a stronger purifying selection of this protein in this group (Table 3). Meanwhile, Laurasiatheria yielded a significantly higher foreground ω and evidence of episodic positive selection, identifying 4 PSS in this group (Table 3), but only one of these residues was identified under pervasive positive selection in previous analysis. In case of CRISP2, primates were the sole group showing a significant acceleration of evolutionary rate (Table 3), suggesting a major influence of positive selection driving CRSP2 evolution in this group. Nonetheless, branch-site model allowing selection was not significant for Primates, nor any other mammalian group (Table 3). For CRISP3, no mammalian group showed an evolutionary rate significantly deviating from the background, and a weak signal of positive selection, with a single PSS identified, was found in Laurasiatheria (Table 3).

**Table 3.** Parameter estimates of branch models to detect selection in mammalian groups

| Gene | Group | Lineages | $\omega_f^a$ | $\omega_b^a$ | LRT<sup>b</sup><br>(M2R - M0) | LRT<sup>c</sup><br>(BS - BSfix) | Number of<br>PSS<sup>d</sup> |
| --- | --- | --- | --- | --- | --- | --- | --- |
| CRISP1 | Glires | 10 | 0.28 | 0.32 | 17.4** | 0 | - |
|  | Primates | 11 | 0.52 | 0.43 | 1.56 | 2.03 | - |
|  | Laurasiatheria | 25 | 0.52 | 0.38 | 10.24** | 17.44** | 4 |
| CRISP2 | Glires | 12 | 0.28 | 0.32 | 3.44 | 0 | - |
|  | Primates | 12 | 0.42 | 0.29 | 5.31* | 0 | - |
|  | Laurasiatheria | 18 | 0.3 | 0.3 | 1.36 | 0 | - |
| CRISP3 | Glires | 6 | 0.45 | 0.54 | 2.31 | 0 | - |
|  | Primates | 12 | 0.56 | 0.59 | 0.31 | 0 | - |
|  | Laurasiatheria | 18 | 0.53 | 0.5 | 0 | 11.68** | 1 |

<sup>a</sup>Average omega (dN/dS) of foreground ( $\omega_f$ ) and background ( $\omega_b$ ) branches.
<sup>b</sup>Likelihood Ratio Tests comparing two-ratio (M2R) and one-ratio (M0) models.
<sup>c</sup>Likelihood Ratio Tests comparing standard branch-site (BS) model with that assuming fixed to 1 (BSfix).
<sup>d</sup>Positively selected sites with Posterior Probability > 0.95 under significant BS model.

Altogether, these results indicate that selective pressures on CRISPs were heterogeneous across mammalian phylogeny. Nonetheless, the lack of, or low signal of positive selection detected in mammalian groups suggests that CRISPs evolved under a major influence of pervasive positive selection acting on the same sites across all lineages, rather than by episodic selection focused on specific lineages.

## Conclusions

In the present study, we reassessed the molecular adaptation of mammalian CRISPs, a family of reproduction-related proteins. Our results revealed evidence of positive selection driving the evolution of the three principal members of the mammalian CRISP family. These findings are consistent with the general assumption that proteins involved in sexual reproduction often evolve adaptively (Swanson and Vacquier 2002; McDonough et al. 2016). Given the involvement of mammalian CRISPs in sperm function and gamete interactions, rapid changes in these proteins could be associated with differences in male reproductive success and with post-copulatory conflicting interests of males and females. Integration of evolutionary and biochemical approaches comparing closely related species of different mating systems will provide meaningful insights into the contribution of mammalian CRISPs to male fertility, sexual competitiveness, and reproductive isolation.

## Materials and Methods

Coding sequences were retrieved from the genomic databases at NCBI (https://www.ncbi.nlm.nih.gov/) and Ensembl (http://www.ensembl.org/). The compiled CRISP sequences (with the accession numbers and organisms) are listed in supplementary file S1. Coding sequences were aligned using the MAFFT program (Katoh and Standley 2013), and the multiple sequence alignment were manually curated using AliView (Larsson 2014). Phylogenetic trees were inferred by ML method using PhyML (Guindon et al. 2010) after selecting the best nucleotide substitution model with jModelTest 2 (Darriba et al. 2012). Input alignments and trees used in selection analysis are available in Supplementary Material (files S2-S9).

Codon-based evolutionary analyses were run with Codeml, implemented in PAML v 4.9c (Yang 2007). The M0 model was used to estimate the global ω of the alignments. To detect lineage-specific changes in selective pressures at individual sites, two-ratio (Yang and Nielsen 1998) and branch-site models (Zhang et al. 2005) were applied. To identify amino acid sites undergoing pervasive adaptive evolution, we applied site-specific models (Nielsen and Yang 1998). Three pairs of nested site models were compared to test for positive selection: M2a (selection) with M1a (nearly neutral), and M8 (beta, selection), with M7 (beta) and M8a (beta, neutral). Those sites with an Empirical Bayes (EB) posterior probability > 0.95 were considered to have evolved under positive selection (Yang et al. 2005). Likelihood Ratio Tests (LRT) were used to compare null and alternative models, approximating the statistic to a chi-squared distribution. Additional likelihood methods to detect pervasive selection analysis (FEL, REL) were applied with Datamonkey 2.0 server (Weaver et al. 2018).

Three-dimensional models of mammalian CRISPs were obtained by using the Swiss-Model server (Arnold et al. 2006). Templates were searched using human CRISP sequences (UniProt ref: P54107, P16562, P54108). The crystal structure of the Stecrisp from the snake *Trimeresurus stejnegeri* (PDB: 1RC91), which met the highest sequence similarity and lowest resolution, was chosen as the template for building the structural models of mammalian CRISPs.

## Acknowledgments

The authors would like to thank Jose Luis De la Vega, Paulina Torres, Francisco Herrera and Shirley Ainsworth for technical assistance; Juan Manuel Hurtado, Roberto Rodríguez, David Santiago Castañeda, Omar Arriaga and Arturo Ocádiz for computing services; and Miguel Arenas for the critical revision of this manuscript. Computational analyses were supported by the cluster of the Instituto de Biotecnología UNAM (http://teopanzolco.ibt.unam.mx//).

